# Effect of Circadian Rhythm Modulated Blood Flow on Nanoparticle based Targeted Drug Delivery in Virtual *In Vivo* Arterial Geometries

**DOI:** 10.1101/2024.06.05.597680

**Authors:** Shoaib A. Goraya, Shengzhe Ding, Mariam K. Arif, Hyunjoon Kong, Arif Masud

## Abstract

Delivery of drug using nanocarriers tethered with vasculature-targeting epitopes aims to maximize the therapeutic efficacy of the drug while minimizing the drug side effects. Circadian rhythm which is governed by the central nervous system has implications for targeted drug delivery due to sleep-wake cycle changes in blood flow dynamics. This paper presents an advanced fluid dynamics modeling method that is based on viscous incompressible shear-rate fluid (blood) coupled with an advection-diffusion equation to simulate the formation of drug concentration gradients in the blood stream and buildup of concentration at the targeted site. The method is equipped with an experimentally calibrated nanoparticle-endothelial cell adhesion model that employs Robin boundary conditions to describe nanoparticle retention based on probability of adhesion, a friction model accounting for surface roughness of endothelial cell layer, and a dispersion model based on Taylor-Aris expression for effective diffusion in the boundary layer. The computational model is first experimentally validated and then tested on engineered bifurcating arterial systems where impedance boundary conditions are applied at the outflow to account for the downstream resistance at each outlet. It is then applied to a virtual geometric model of an *in vivo* arterial tree developed through MRI-based image processing techniques. These simulations highlight the potential of the computational model for drug transport, adhesion, and retention at multiple sites in virtual *in vivo* models. The model provides a virtual platform for exploring circadian rhythm modulated blood flow for targeted drug delivery while minimizing the *in vivo* experimentation.

**Statement of Significance:** A novel integration of nanoparticle-based drug delivery framework with shear-rate dependent blood flow model is presented. The framework is comprised of a unique combination of mechanics-based dispersion model, an asperity model for endothelium surface roughness, and a stochastic nanoparticle-endothelial cell adhesion model. Simulations of MRI based *in vivo* carotid artery system showcase the effects of vessel geometry on nanoparticle adhesion and retention at the targeted site. Vessel geometry and target site location impact nanoparticle adhesion; curved and bifurcating regions favor local accumulation of drug. It is also shown that aligning drug administration with circadian rhythm and sleep cycle can enhance the efficacy of drug delivery processes. These simulations highlight the potential of the computational modeling for exploring circadian rhythm modulated blood flow for targeted drug delivery while minimizing the *in vivo* experimentation.

## 1. Introduction

Diseases of the central nervous system (CNS) are among the leading causes of death in the United States, with Alzheimer’s disease alone being responsible for 121,499 deaths in 2019 [1]. Even in non-fatal cases, these CNS conditions pose a tremendous health burden [2]. Treatment of these diseases is limited in part by the difficulty of delivering therapeutics to the site of the diseased tissue due to the blood–brain barrier (BBB) [3]. Extensive efforts have been dedicated to optimizing the delivery of therapeutic drugs of interest to diseased tissues through the bloodstream while minimizing unwanted interaction with healthy tissues and subsequent adverse reactions [4]. One prominent approach to overcome this barrier is to encapsulate drug molecules within biodegradable nanoparticles tethered with bioactive ligands capable of binding to receptors that are overexpressed in the microvascular networks of target tissues [5]. The mechanism of the transport across the BBB appears to be receptor-mediated endocytosis followed by transcytosis of the drug-loaded nanoparticles into the brain or by the release of the drugs within the endothelial cells [5–8]. Nanoparticle surface treatment is required in order to enable this receptor-mediated uptake [5–7]. Numerous studies in animal models, as well as clinical investigations, have provided evidence that the pharmacokinetics and/or the efficacy of drugs can be enhanced by the circadian rhythm, in addition to the timing of drug administration within the 24-hours cycle of a day [9]. However, little attention has been paid to the influence of chronobiology on the processes of nanoparticle transport into the brain, and subsequent retention at the target sites [10].

The circadian rhythm governs the 24-hour cycle of bodily changes impacting sleep, metabolism, and blood flow in response to natural cues. Disruptions to this cycle correlate with adverse health outcomes like increased risk of myocardial infarction/ stroke and worsened eczema severity due to sleep disruptions [11, 12]. Circadian rhythms also affect drug metabolism enzymes, with implications for targeted drug delivery, especially in chronotherapy [13, 14]. Chronotherapy involves administering drugs at specific times to optimize effectiveness and minimize side effects based on circadian cycles and physiological changes [14]. Circadian rhythms influence the pharmacokinetics and pharmacodynamics of medications, suggesting potential improvements in therapeutic impact by aligning drug delivery with circadian regulation [15]. Synchronizing targeted drug delivery with the circadian clock can enhance efficacy and enable personalized medicine, reducing the risk of adverse effects. Peptide-ligated nanoparticle drug delivery systems, responsive to circadian cues, offer a means to release drugs at optimal times, improving therapeutic impact.

Several microfluidic devices have been proposed for *in vitro* studies of nanoparticle transport, aiming to enhance understanding in the *in vivo* microvasculature [3, 16–18]. However, these models lack insights into the impact of circadian rhythm on particle transport, exhibit limitations in predicting drug efficacy for patient-specific attributes and require substantial resources [19]. Additionally, the crucial aspect of particle detachment in external flow has been overlooked in prior studies [20–22]. The retention of attached particles plays an important role in determining the efficacy of drug delivery systems, as the concentration of attached particles at the target site diminishes over time due to dislodging hemodynamic forces [23].

With the objective to investigate the intricate interactions between nanoparticles and biological systems, we present a computational model for nanoparticle-facilitated drug delivery to determine factors that affect particle transport and retention at targeted sites under circadian blood flow fluctuations. PLGA-*b*-HA nanoparticles decorated with VHSPNKK peptides are represented as a homogenized mixture in the fluid and advection-diffusion equation is used to simulate the formation of drug concentration gradients in the blood flow. A novel integration of Taylor-Aris dispersion model [24, 25] and an asperity model with Decuzzi’s particle-cell adhesion model [26] is presented to predict nanocarrier transport, attachment, and detachment more accurately. The dispersion model accounts for shear-induced diffusion of particles in the boundary layer, while the asperity model accounts for the interaction between the fluid and endothelial cell layer. The computational method offers a distinctive approach by integrating mechanics-based drug delivery models into blood flow modeling approaches in patient-specific geometries.

The drug delivery model is implemented in a stabilized finite element framework [27–29] to accurately capture concentration gradients in convection-dominated flow conditions. Blood rheology is accounted for via a shear-rate-dependent model [30, 31]. Circulatory effects downstream of the modeled arterial system are incorporated using resistance outflow boundary conditions. Physiologically relevant pulsatile inflow rates and pressure profiles are used to determine resistance functions. Circadian variation in hemodynamics is modeled using clinical data [32, 33]. Particle-cell adhesion model is embedded via Robin boundary conditions (BC) applied at the endothelial cell layer surface [34, 35]. Robin BC combines Neumann (drug flux in normal direction) and Dirichlet (drug concentration) boundary conditions through the vascular deposition parameter [20]. This parameter fully accounts for particle-cell binding kinetics by integrating information of biochemical, biophysical, and geometric properties from micro/ nano scale into macro scale.

The proposed method was validated through carefully designed experiments. *In vitro* experiments were carried out in a 3D microfluidic chamber, lined with TNF-*α* treated mouse endothelial cells with enhanced VCAM expression, to determine particle adhesion and retention at the target site. Atomic force microscopy (AFM) was used to determine the binding affinity between coated nanoparticles and the ligand-coated receptor cells. This data was used to determine the biochemical parameters in the probabilistic particle-cell adhesion model. The validated model was then employed to simulate the effect of circadian modulation of blood flow on targeted drug delivery via nanoparticles in virtual *in vivo* geometries.

An outline of the rest of the paper is as follows: Section 2 presents mechanics-based models that are operational at multiple scales and are embedded in the mathematical framework for targeted drug delivery. A stabilized finite element method is presented in Section 2.7. Discussion on circadian rhythm in human body and its impact on blood flow dynamics in presented in Section 2.8. Experimental validation of the computational method via an *in vitro* microfluidic device is shown in Section 3. Numerical test cases of idealized bifurcating artery, idealized arterial tree, and a patient-specific carotid artery system are presented in Section 3. These simulations showcase the application of computational techniques for targeted drug delivery in virtual *in vivo* models and the impact of circadian rhythm on drug transport and adhesion. Conclusions are drawn in Section 4.

## 2. Materials and Methods

### 2.1 Governing equations for targeted drug delivery in vasculature

The balance laws for transport of ligand-coated nanoparticles in an incompressible shear-rate dependent fluid (blood) are given below:

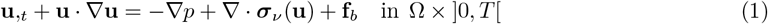

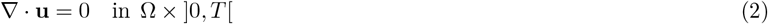

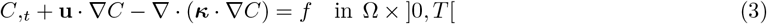

where **u** (**x**, *t*) and *p* (**x**, *t*) are the velocity and kinematic pressure fields, respectively; *C* (**x**, *t*) is the volume concentration of particles; ***σ***_*ν*_ is the deviatoric stress tensor which in the case of shear-rate dependent fluids depends on the viscosity of the fluid, and *κ* is the diffusivity tensor; **f**_*b*_ is the body force for the mechanical field, and *f* is the source/sink term for the concentration field. Equation (1) is the momentum balance equation, equ. (2) is the continuity equation for conservation of mass and it enforces the incompressibility constraint, and equ. (3) is the convection-diffusion equation for the evolution of particle concentration field.

The mathematical model is supplemented with the initial conditions in the domain **x**∈Ω at time *t* = *t*_0_ and boundary conditions on the domain boundary G,

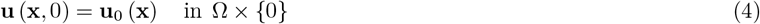

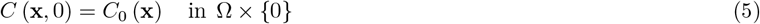

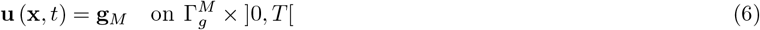

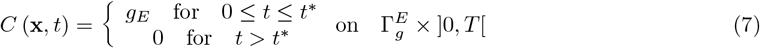

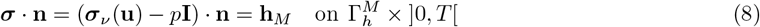

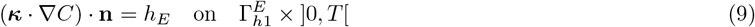

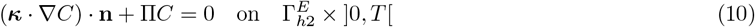

where **u**_0_ and *C*_0_ are the initial conditions for velocity and concentration fields, respectively; **g**_*M*_ and *g*_*E*_ are the prescribed values for velocity and concentration fields, respectively; while **h**_*M*_ and *h*_*E*_ are the prescribed fluxes for the velocity and concentration fields, respectively. *t*^∗^ denotes the time duration for the injection of particles at the inlet. ***σ*** is the total stress in the fluid and **n** is the outward normal vector at the boundary.

The boundaries satisfy the following conditions: 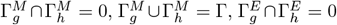 and 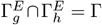. Equation (10) defines Robin boundary conditions that were used for particle adhesion to the endothelial cell layer. Π is the positive-valued adhesion factor that depends on particle-cell adhesion model [26, 36].

### 2.2 Shear-rate dependent blood flow model

In this work we have employed shear-rate dependent non-Newtonian model for blood [37]. The flow of shear-thinning fluids generally gives rise to pseudo-plastic velocity profiles that are characterized by lower velocities due to low convective flow but sharper and thinner boundary layers [27]. The ability to capture the boundary layer is of significant importance for shear-thinning flows. Consequently, the stabilized methods for this class of fluids were developed by the senior author, and the interested reader is referred to [30, 31, 38, 39].

The Cauchy stress tensor is split into volumetric stress and viscous stress components by treating pressure as an independent field:

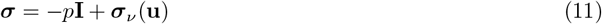

where ***σ*** is Cauchy stress tensor, *p* is the pressure field or the volumetric stress, and ***σ***_*ν*_ (**u**) is the nonlinear viscous stress given as:

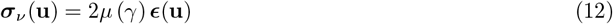

where ***ϵ***(**u**) is the rate-of-deformation tensor which is defined as ***ϵ*** (**u**) = 1*/*2 ∇**u** + (∇**u** ^*T*^). In shear-rate dependent fluids, the viscosity field *µ* (*γ*) is a nonlinear function of the shear-rate *γ* defined as 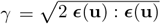

In this work we have employed the Carreau-Yasuda model [37] to represent the shear-thinning behavior of blood. In shear-rate dependent fluids, the nonlinear function of the viscosity is given by:

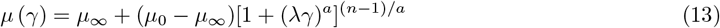

where *µ*_0_ and *µ*_∞_ are asymptotic viscosities at zero and infinite shear-rate, respectively, and *a, n* and *λ* are empirically determined constitutive parameters. Coefficients *a* and *n* are non-dimensional parameters that control the shear-thinning or shear-thickening behavior of fluids in the non-Newtonian regime between the two asymptotic viscosities. The Carreau–Yasuda model possesses constant values of viscosity at both low and high ranges of shear-rate, and a varying viscosity in the intermediate range of shear-rate. The model reverts to the Newtonian fluid model by setting *µ*_0_ = *µ*_∞_ as shown in Figure 2.

#### Remark 1.

The computational model presented here is general and any constitutive model for the shear-rate dependent response of the fluid can be incorporated in it.

### 2.3 Resistance boundary conditions

The cardiovascular system is comprised of heart and blood vessels that form a closed network. Developing a computational model for the entire system is not only difficult due to its geometric complexity, but it is also computationally expensive. Consequently, developing appropriate outflow boundary conditions that can take into account the effects of the circulation system downstream of the zone of interest is an essential ingredient of blood flow modeling strategies. With appropriate boundary conditions applied at the outflow surfaces, blood flow simulations can be carried out only in the network of interest, and this can substantially reduce the cost of computation while preserving the accuracy of the modeled physics.

The downstream resistance *R* can be assumed to have a linear relation between the flow-rate and the pressure. The mean pressures and flow-rates at outlets determine the constant resistance values at these outlets.

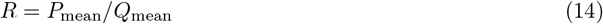

where *P*_mean_ and *Q*_mean_ are the mean pressure and flow-rate for one generic cardiac cycle at the outflow boundary, respectively. The resistance boundary conditions that produce physiological pressure waveform at the outflows are imposed at all the outlets G_out_ to incorporate the downstream resistive effects of the blood vessels. It is given as:

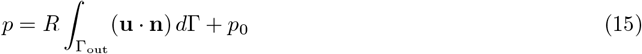

where *p*_0_ is the constant downstream pressure. This method also allows for embedding clinically measured downstream resistance via a functional form for the outflow boundary conditions. In this case flow-rate and pressure profiles of a typical carotid artery are used to determine time-dependent resistance functions.

### 2.4 Particle-cell adhesion model

The ability of the drug coated nanoparticles to adhere to the target endothelial cells is a critical step in targeted drug delivery processes. Since we have adopted a continuum modeling approach, information at the micro- and nanoscale is embedded in the computational model via the Robin boundary condition that is applied at the targeted vessel wall G_wall_.

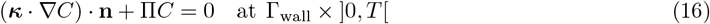

where Π = *P*_*a*_*γ* (*d*_*p*_*/*2) is vascular deposition parameter, *γ* is the shear rate defined in Section 2.2, *d*_*p*_ is the diameter of nanoparticles, and *P*_*a*_ is the probability of adhesion. The Robin boundary condition accounts for particle-cell adhesion based on the probability of adhesion *P*_*a*_ [26, 36]. *This probability is directly related to the strength of adhesion: the larger is the value of P*_*a*_ the larger is the adhesive strength of the particle to the endothelial cell layer. Only the particles in the boundary layer of the fluid filled vessel experience the binding affinity (Figure 1). Thus equ. (16) integrates information from the macroscale (vessel geometry and flow conditions) with data from the micro- and nanoscale (particle geometry, receptor density, and affinity), thereby avoiding massive, computationally inefficient discretization over multiple spatial and temporal scales. Moreover, the mass flux of particles in the direction normal to the wall can be related to the local increase in mass of particles adhering per unit surface *φ* (**x**, *t*):

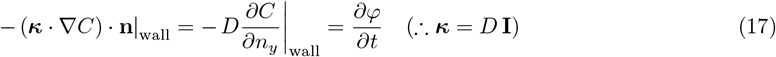

The probability of adhesion *P*_*a*_ includes 2D binding characteristics of receptors and ligand coated particles that are attached to surfaces, and their bonding is subjected to dislodging hydrodynamic forces. An experimentally validated closed form expression for the probability of adhesion is given in [26, 36]. The expression describes cellular adhesion by coupling the bond forces to the binding affinity. The probability of adhesion for ellipsoid particles is given as [26]:

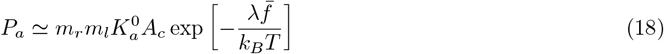

where 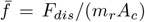 is the force per unit ligand-receptor pair, *A*_*c*_ is the area of interaction between ligand and receptor, *F*_*dis*_ is the dislodging force due to drag force and rotating torque, 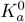 is ligand-receptor affinity constant at zero load, *m*_*l*_ is surface density of ligand molecules (#/*m*^2^), and *m*_*r*_ is surface density of receptor molecules (#/*m*^2^). The dislodging force *F*_*dis*_ also includes the effect of equilibrium separation distance between the cell layer and the particle, and the maximum distance at which ligand–receptor bonds can be formed [26]. The above expression shows that the strength of adhesion is affected by the geometric features of the particle (particle size and aspect ratio), by the biophysical parameters (dislodging force depending on the strength of ligand–receptor bonds and wall shear stress, and the surface densities of the ligands *m*_*l*_ and of the receptors *m*_*r*_), and by the biochemical parameters (characteristic length *λ* of the ligand–receptor bond and the characteristic affinity constant 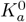 of the ligand–receptor pair).

**Figure 1:**
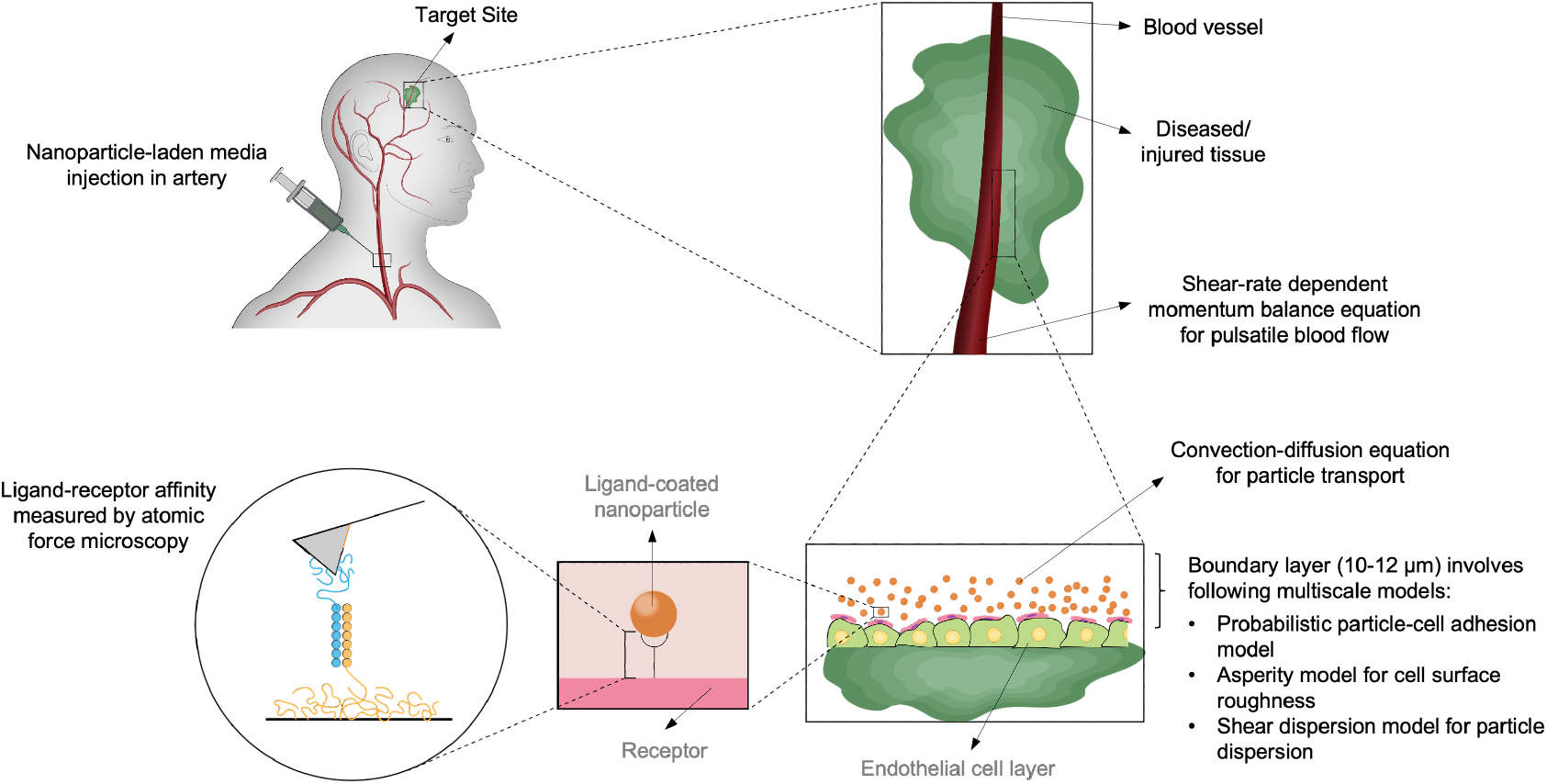
Schematic diagram of drug delivery using nanocarriers designed for active targeting in circulation. The experimentally validated computational model serves as a digital twin, enabling investigation of the effects of hemodynamic forces and particle geometry on particle adhesion in patient-specific arterial models. A shear-rate dependent blood flow model, along with resistance boundary conditions for downstream pressure, is used to simulate pulsatile blood flow in the arterial system. Adhesion kinetics are simulated through various multiscale models, incorporating experimentally measured parameters, within the boundary layer at the target sites.

**Figure 2:**
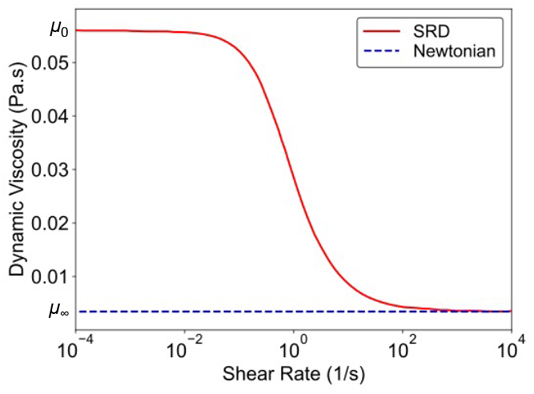
Viscosity of Newtonian and shear rate dependent (Carreau-Yasuda) constitutive models for blood. The Carreau-Yasuda model for blood shows changes in viscosity as a function of the shear-rate, while the Newtonian model has a constant viscosity at all shear-rates.

For spherical particles, equ. (18) can be simplified to:

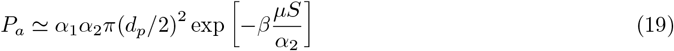

where the parameters *α*_1_, *α*_2_, and *β* are given as:

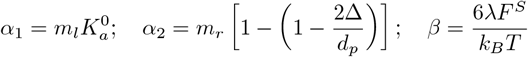

In the above parameters, Δ is the distance between particles and endothelial cell layer at equilibrium, *λ* is ligand-receptor bond length, and *F*^*S*^ is drag coefficient that depends on the particle geometry.

Particle-cell binding is an interrelation of random processes at distinct time scales. Particles undergo rapid binding and transport, resulting in a stochastic spatial distribution of bound particles fluctuating about some mean distribution. Moreover, it was also observed during the experiments [23] that cells are clustered together at random locations giving rise to non-uniform distribution of receptor density. To incorporate this effect into our mathematical framework, we have employed Gaussian sampling of receptor densities over the endothelial cell layer. Gaussian sampling was done from a normal distribution with a coefficient of variation of 0.1. This resulted in a more realistic picture of the particle-cell binding process which is stochastic in nature. The resulting adhesion model when integrated in the computational framework provides a mechanistic understanding of the magnitude and distribution of nanoparticles that adhere to the endothelial layer.

The parameters to compute *P*_*a*_ in the adhesion model were obtained through characterization experiments that are reported in Table 1. Atomic force microscopy (AFM) was used to determine the bond-length and the equilibrium distance between the peptide and the protein. Meanwhile, the ligand and receptor densities, as well as the ligand-receptor affinity constant, were adopted from the published literature [20, 36]. The binding affinity was observed to decrease in experiments where particles were not treated with VHSPNKK peptides. This decrease was accounted for in the adhesion model by calibrating the binding affinity constant with the experimental data.

**Table 1:**
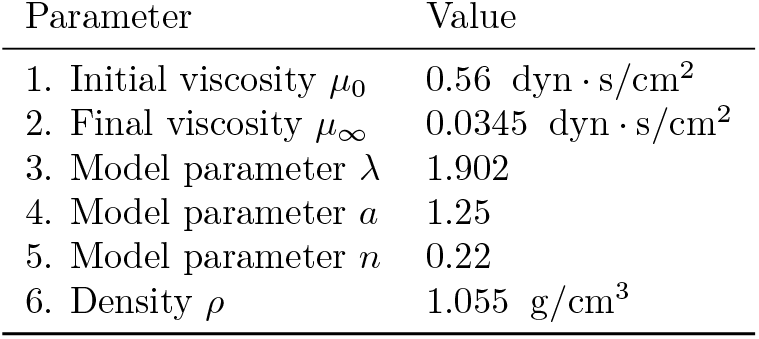
Parameters for Carreu-Yasuda blood flow model.

| Parameter | Value |
| --- | --- |
| 1. Initial viscosity $\mu_0$ | 0.56 dyn · s/cm<sup>2</sup> |
| 2. Final viscosity $\mu_\infty$ | 0.0345 dyn · s/cm<sup>2</sup> |
| 3. Model parameter $\lambda$ | 1.902 |
| 4. Model parameter $a$ | 1.25 |
| 5. Model parameter $n$ | 0.22 |
| 6. Density $\rho$ | 1.055 g/cm<sup>3</sup> |

### 2.5 Modeling the surface roughness of endothelial cell layer

Modeling of particle transport and adhesion requires taking into account the effect of interaction between fluid in the boundary layer with the irregular surface of the endothelial cell layer. The irregular surface induces roughness that helps in the retention of the attached particles. In numerical test cases, an idealized smooth cell surface failed to produce experimentally observed particle retention during the detachment/ wash flows. Accordingly, we employed a two-body contact formulation to model the particle-cell interaction. The mathematical formulation of the two-body contact problem involves constraint equations, known as the Kuhn-Tucker conditions, to describe the relationship between the contact pressure *q*_*N*_ and penetration or gap **g**_*N*_ in the surface normal direction [40]. In addition, coupling in the tangential direction is expressed through conjugate variables, the tangential gap **g**_*T*_ and the tangential flux **q**_*T*_. The gap functions are defined as [41]:

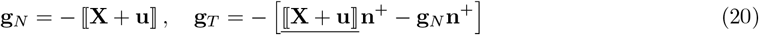

where ⟦· ⟧ and 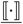 are jump operators along the interface of the two bodies with dot product and tensor product, respectively. We make use of the fact that the fluid and cell layer are always in contact with each other, thereby rendering the relative position vector **X** = **0**. Moreover, the cell layer does not move and has zero velocity. This simplifies (20) to:

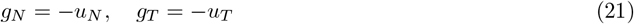

where *u*_*N*_ and *u*_*T*_ are normal and tangential components of the fluid velocity, respectively. Embedding Kuhn-Tucker contact conditions into convection-diffusion equ. (3) and ignoring the normal component *g*_*N*_ yields the following:

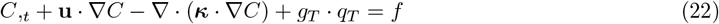

Numerical modeling of asperities as shown in Figure 1 requires discretizations that are at least an order of magnitude smaller than the asperity size. Since endothelial cells are in the range of 0.1 *µ*m, therefore it is appropriate to use micromechanics asperity model for contact between fluid and irregular cell layer surface. We employed the well-known Coulomb friction model at the mean asperity height to model surface roughness due to irregular endothelial cell layer. Coulomb model couples the normal and tangential tractions with a proportionality constant called the friction coefficient *µ*_*f*_. The tangential flux given by the Coulomb model is:

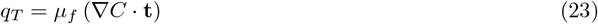

where **t** is tangential vector at the surface of the cell layer. The friction term in equ. (22) was only activated at a mean asperity height *d* from the bottom of the vessel (Figure 1). The friction coefficient between fluid and endothelial cell layer was estimated empirically by calibrating the model equ. (23) with the experimental data.

### 2.6 Dispersion of particles in the boundary layer

In confined flows, dispersion due to shear (velocity gradient) affects the mixing and transport of the particles in the boundary layer region (Figure 1). Dispersion increases the dislodging hydrodynamic forces which impacts the number of nanoparticles that are retained at the endothelial cell layer during the washing stage. This effect is incorporated by using Taylor-Aris expression for effective diffusion in the boundary layer region that takes into account longitudinal dispersion [24, 25]:

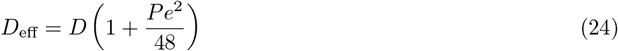

where *Pe* = *UL*/*D* is the Peclet number, *U* is the mean flow velocity, and *L* is the length scale associated with the problem. The length scale chosen is diameter of the particle *d*_*p*_. *D* is isotropic particle diffusivity coefficient obtained from Stokes-Einstein equation *D* = *k*_*B*_*T* /(3*µd*_*p*_) where *k*_*B*_*T* is the Boltzmann thermal energy (4.142 *×* 10^−21^ J) and *µ* is the dynamic viscosity of the fluid.

### 2.7 Stabilized finite element method

In this section we present the synopsis of the stabilized finite element method that provides a fully coupled system of equations that can be consistently linearized to develop a monolithic solution procedure that yields quadratic rate of convergence in the nonlinear solution strategies.

#### 2.7.1 Standard weak form

The standard weak form of the problem is: Find **V** = (**u**, *p, C*) ∈ 𝒰_*t*_ *×*𝒫_*t*_ *×*𝒞_*t*_, such that ∀ **W** = (**w**, *q, η*) ∈ 𝒲 *×* 𝒬 *×* ℋ

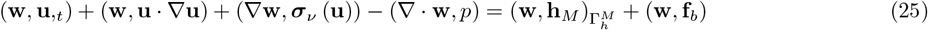

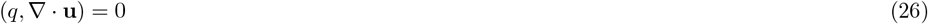

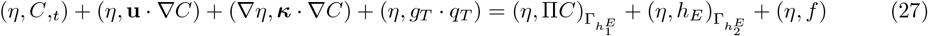

where 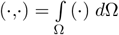 denotes the *L*_2_ (Ω) inner product.

The appropriate functional spaces for the velocity, pressure, and concentration trial solutions, 𝒰_*t*_, 𝒫_*t*_, 𝒞_*t*_ are defined below:

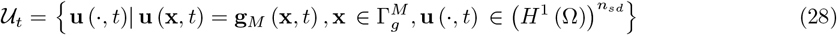

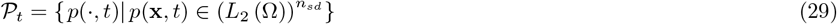

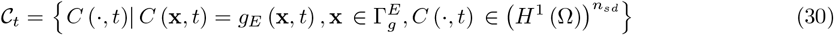

where *L*_2_ (Ω) and *H*^1^ (Ω) are the standard Sobolev spaces. Let 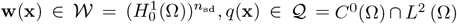, and 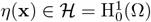 be the weighting functions for the velocity **u**, kinematic pressure *p*, and concentration *C*, respectively. The weighting function spaces 𝒲, 𝒬, ℋ are the corresponding spaces of equ. (28)-(30) that satisfy the homogeneous part of the essential boundary conditions.

#### 2.7.2 Variational multiscale method

The bounded domain Ω is discretized into non-overlapping subdomains 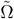 with subdomain boundary 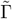, such that 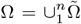 and *n* is the total number of subdomains that comprise Ω. The union of interior of subdomains is denoted as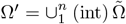, and the union of subdomain boundaries is 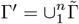. Using the scale split feature of the variational multiscale method [42, 43] for momentum balance equation [44] and advection equation [45], we decompose the trial solution fields (**u**, *C*) into coarse and fine scales. In the context of nonlinear problems, this decomposition is to be viewed as a projection of sub-grid scales onto resolved scales, e.g., for the velocity field **u**^*′*^ = P(**u** − ū).

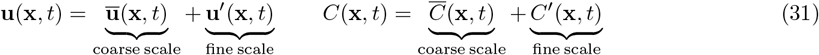

The spaces of coarse scale functions are linearly independent of that of the fine scales, i.e., 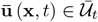 and 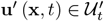 where 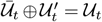 and 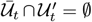. The same holds for the concentration field. Similarly, the corresponding weighting functions (**w**, *η*) are decomposed as:

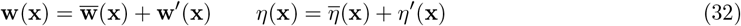

By the additive nature of the decomposition of weighting function in equ. (32), we can split the weak form into coarse-scale and fine-scale sub-problems by grouping the terms depending on the weighting functions at either scale:

##### Coarse-scale sub-problem

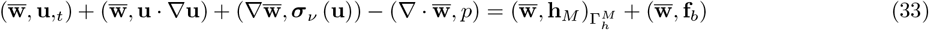

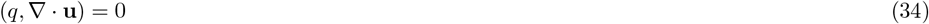

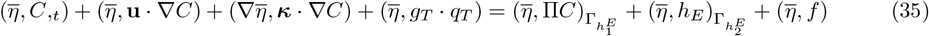

##### Fine-scale sub-problem

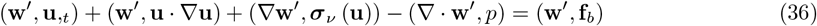

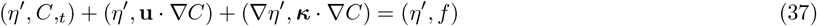

#### 2.7.3 Derivation of fine scale models for velocity and concentration fields

The key idea at this point is to resolve equ. (36) and (37) locally and extract the models for the time-dependent fine-scale velocity and concentration fields, via the residuals of the Euler-Lagrange equations of the coarse-scales, respectively. These models when embedded in the coarse scale formulation provide stability in the sense of *inf-sup* condition as well as high advection velocity. These subgrid models also project missing physics onto the resolved scales. This step restores stability of the mixed weak form and also increase the accuracy of the formulation. The fine-scale sub-problem presented in equ. (36) and (37) yields a nonlinear coupled system. Therefore, we first linearize the fine-scale variational equations with respect to the fine-scale velocity and concentration fields as follows:

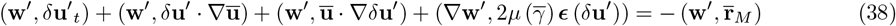

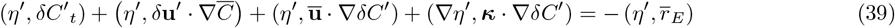

where 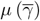 is the nonlinear viscosity that is a function of shear-rate which is calculated based on the coarse-scale velocity field, 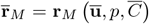 is the residual of the Euler-Lagrange equations of the coarse-scale conservation of momentum, and 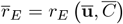 is the residual of the Euler-Lagrange equation of the coarse-scale concentration equation. These residuals are defined as,

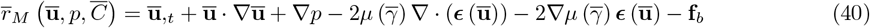

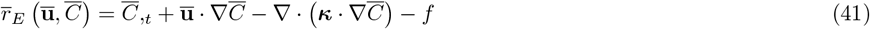

In order to resolve the fine-scale sub-problems (36) and (37), and to derive fine-scale models for the velocity and concentration fields, we make some simplifying assumptions to localize the problems. We assume that the fine-scale trial solutions vanish at the subdomain boundaries, namely, **u**^*′*^ = **0** on **G**^*′*^ and *C*^*′*^ = 0 on G^*′*^. We follow along the lines of [43] and employ bubble functions 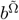 defined over 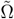 to interpolate the fine-scale trial solutions and weighting functions as follows.

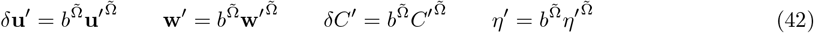

We substitute the above defined fine-scale trial solutions and weighting functions into equ. (40) and (41), apply mean projection theorem, and utilize backward Euler time marching scheme to discretize-in-time the linearized weak forms of equ. (40) and (41). We now resolve the fine-scale problem, and construct the projection from the residual of the coarse-scale Euler-Lagrange equations to the fine-scale solution fields **V**^*′*^ = *{***u**^*′*^, **C**^*′*^*}*. This provides fine-scale solution in terms of the residuals of the Euler-Lagrange equations scaled by the stabilization tensor.

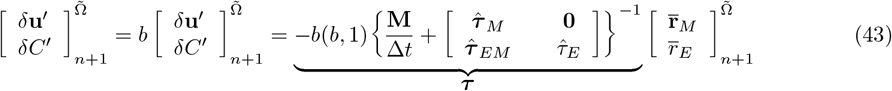

where **M** is the mass matrix and Δ*t* is the time step. For ease of notation, we set subdomain based bubble functions 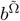 as *b* and remove the superscript 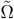 in the subsequent formulation. The explicit form of various terms in equ. (43) are:

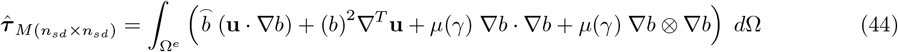

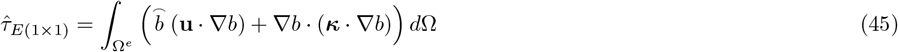

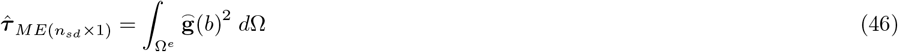

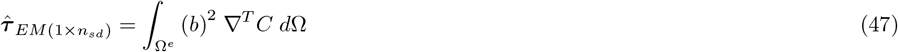

where ***τ*** is the discrete system that relates the fine-scale velocity and concentration fields with the residual of coarse-scale momentum balance and concentration equations. The physical interpretation of 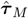 includes the sweeping effect (i.e., the fine-scale velocity transported by the coarse-scale velocity), the distortion effect (i.e., the coarse-scale velocity transported by the coarse-scale velocity), and the fine-scale diffusion; 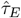 is the discrete system that relates the fine-scale concentration and the residual of the coarse-scale equation of convection-diffusion equation of concentration transport. They are form identical to the discrete fine-scale system of linearized Navier-Stokes equations and a generic advection-diffusion equation from earlier works of the senior author [45–47].

#### 2.7.4 Stabilized formulation

Similarly, we linearize the nonlinear coarse-scale sub-problems in equ. (33)-(35) with respect to the fine-scale velocity and concentration fields, combine the resulting formulation and group the terms that depend on the fine scale fields.

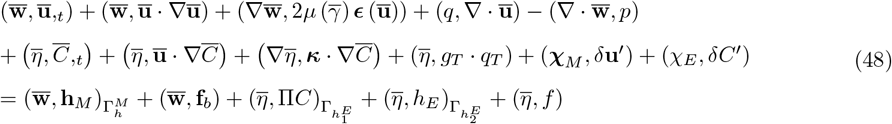

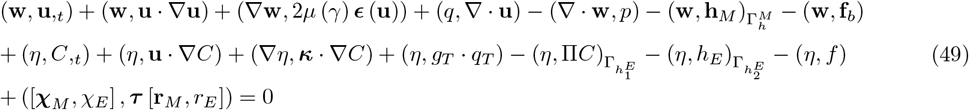

where,

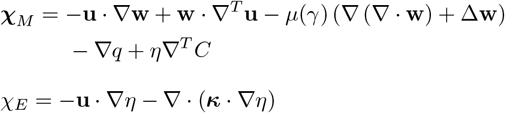

##### Remark 2.

Equation (48) is fully represented in terms of the coarse-scale fields, therefore, we drop the superposed bar from the coarse-scale weighting function and trial solutions and perturbation notation *δ* before fine-scale trial solution fields.

By substituting the fine-scale solutions defined in equ. (43) into the corresponding slots in the coarse-scale formulation, we arrive at the stabilized form that is expressed in the residual form as follows. Formal statement is: Find **V** = (**u**, *p, C*) ∈ 𝒰_*t*_ *×* 𝒫_*t*_ *×* 𝒞_*t*_, such that ∀ **W** = (**w**, *q, η*) ∈ 𝒲 *×* 𝒬 *×* ℋ, the following holds.

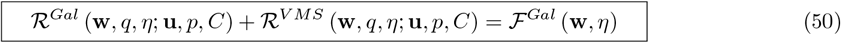

where,

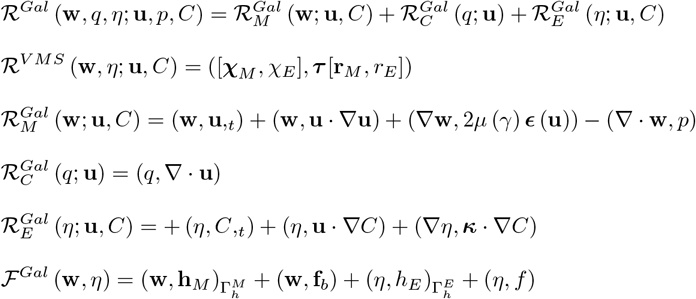

##### Remark 3.

For computational expediency, in the development of the discrete algebraic problem, we set 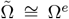 where Ω^*e*^ is the domain of a generic element in the mesh. In this case, 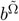 is represented by polynomial bubble *b*^*e*^(*ξ*) defined over Ω^*e*^ such that *b*^*e*^(*ξ*) = 0 on the boundary of Ω^*e*^.

### 2.8 Circadian rhythm modulated blood flow

Circadian rhythm is primarily governed by the body’s intrinsic timing system, the suprachiasmatic nuclei (SCN) located in the hypothalamus of the brain, which works in conjunction with the light/dark cycle to synchronize the wake/sleep cycle and other physiological processes, including blood pressure. Blood pressure typically follows a predictable pattern over the 24-hour circadian cycle due to heart rate (HR) and systemic vascular resistance (SVR) changes that occur due to the effects of peptides, neurotransmitters, and hormones, which are regulated by the SCN [48]. This pattern typically involves a steep rise in systolic blood pressure (SBP) and diastolic blood pressure (DBP) when awakening in the morning with an afternoon decline, followed by a subsequent decline in SBP and DBP in the evening and into sleep [49]. SCN also influences short-term variations in these signaling molecules, causing short-term blood pressure oscillations over the course of the wake/sleep cycle [48].

Cell physiology is also regulated by the SCN and shows rhythm in cellular metabolism and proliferation, with predictable amplitudes and times of peaks and troughs. It can be used to reduce harmful effects of cytotoxic treatments by administering drugs that target the regulation of cell-cycle events or angiogenesis at specific times (chronotherapy). Pre-clinical and clinical evidence supports investigations of the chronotherapy hypothesis in cancer patients [50].

Our objective in this paper has been to investigate how the circadian rhythm influences the transport of drug via blood circulation to specific target sites. Consequently, our focus is revolves around understanding how the circadian cycle impacts the physiological aspects of circulation. Figure 3 shows variations in systolic blood pressure (SBP) and diastolic blood pressure (DBP) in dipper patients who display the typical nocturnal decrease in blood pressure. This decrease in blood pressure correlates with the natural oscillations of heart rate (HR) and systemic vascular resistance (SVR) over the course of the circadian cycle. The clinical data in Figure 3 was obtained on dipper patients in [32, 33].

**Figure 3:**
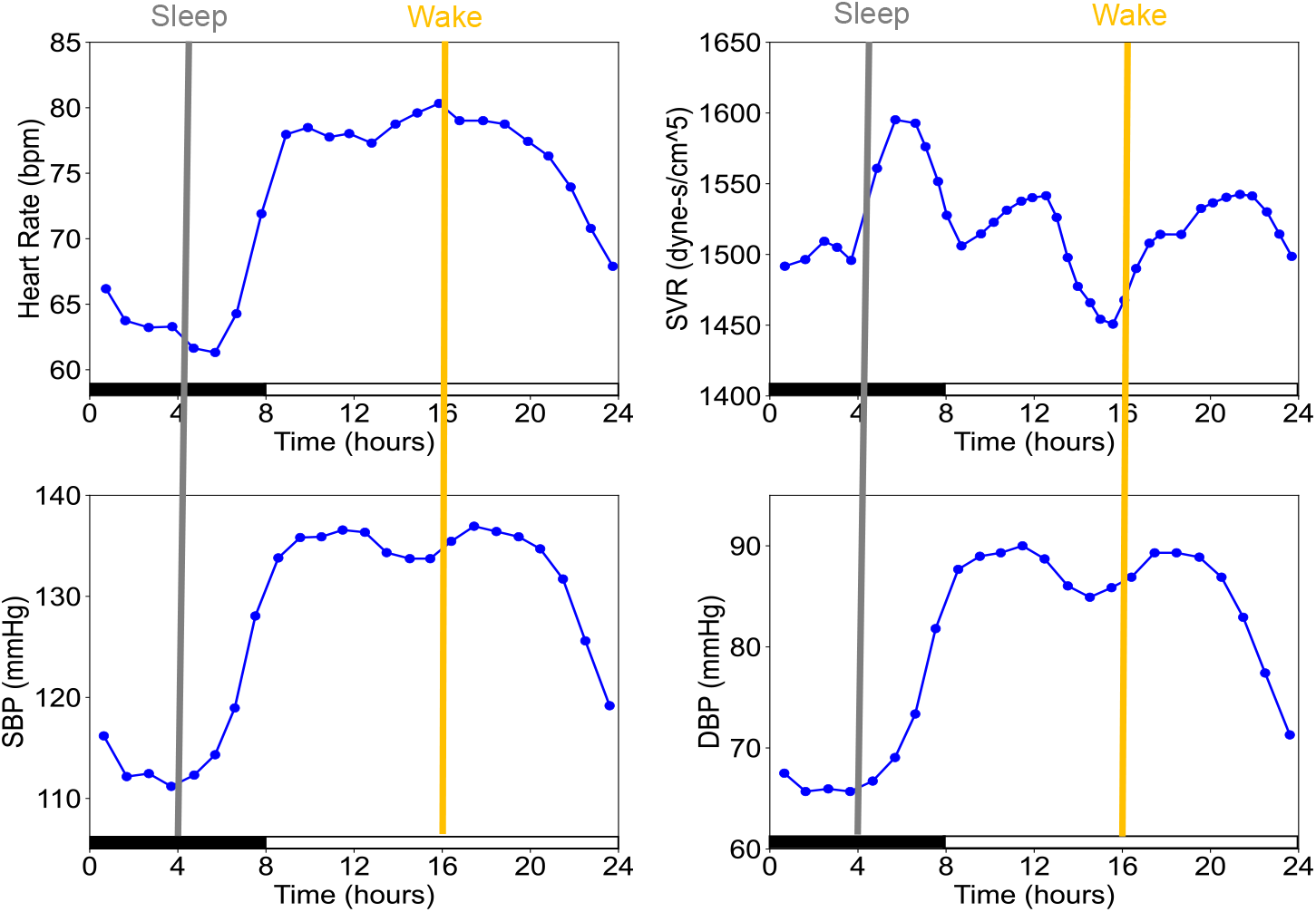
Experimental data of dipper subjects showing 24-hour variation of systolic blood pressure (SBP), heart rate (HR), and diastolic blood pressure (DBP) as reported in [32], and systemic vascular resistance (SVR) as reported in [33]. The data is organized such that the sleep cycle begins at 12 A.M. and ends at 8 A.M. The magenta and yellow lines show parameters at which sleep and wake cycles are simulated, respectively.

## 3. Results and Discussion

We first present experimental validation of the numerical method presented above. In [23], the method was validated on a 3D microfluidic chamber that was lined with C166 mouse endothelium cells at the bottom surface. A solution of PLGA-*b*-HA nanoparticle with water as a base fluid was injected in the chamber to facilitate particle adhesion. In the present paper, we investigate the performance of the model on biologically relevant arterial geometries under *in vivo* flow conditions. Furthermore, the blood is considered as a fluid with shear-rate dependent behavior. The resistance from the distel part of the arterial network is accounted for via resistance boundary conditions applied at the outflow using equ. (15).

The second and third test cases showcase the performance of the method to predict drug transport and adhesion in idealized engineered arteries with bifurcations. Effect of the circadian rhythm modulated blood flow on nanoparticle adhesion and retention is also investigated. The final test case demonstrates the applicability of the method to an MRI-based geometry of the carotid artery system, highlighting the effects of curvatures and bifurcations on particle adhesion at the target sites.

The stabilized finite element method presented in (50) is implemented using equal order, linear and quadratic, tetrahedral and hexahedral elements. The generalized-*α* method is used for time integration. The nonlinear problem is solved using the Newton–Raphson method and the linear system of equations is solved via the GMRES solver with additive Schwarz preconditioner. Because of the mathematically refined representation of the viscosity field (equ. (12)) in the consistent tangent matrix, the nonlinear solver produces quadratic convergence rate. Figure 4 shows velocity waveform in a carotid artery during a typical cardiac cycle that is applied at the inlet. For engineered bifurcating artery and the idealized arterial tree, the velocity magnitude in Figure 4 is reduced to obtain shear rates within a range of 10−250 s^−1^ which is typically observed in microvasculature [51]. The shear rate dependent Carreau-Yasuda model is employed for non-Newtonian behavior of blood flow for which the parameters are given in Table 1. Parameters that are used in the adhesion model are experimentally determined and are given in Table 2.

**Table 2:**
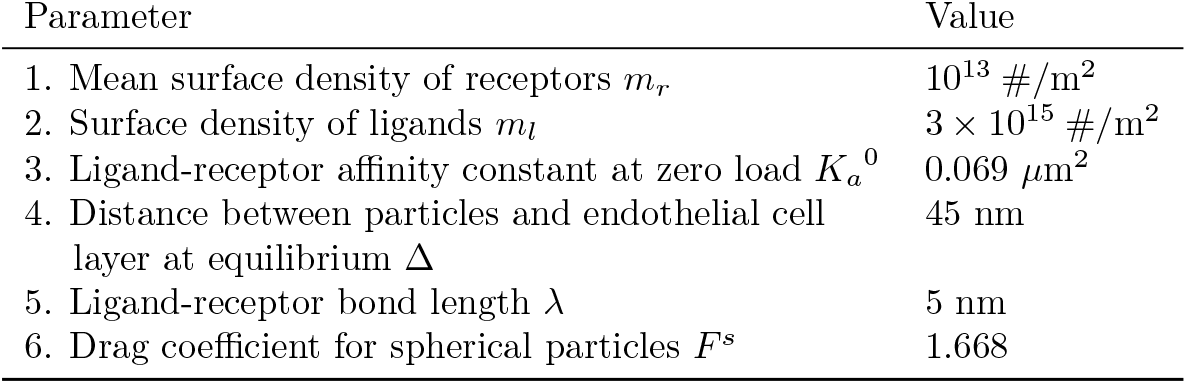
Parameters for particle-cell adhesion model.

| Parameter | Value |
| --- | --- |
| 1. Mean surface density of receptors $m_r$ | 10<sup>13</sup> #/m<sup>2</sup> |
| 2. Surface density of ligands $m_l$ | 3 × 10<sup>15</sup> #/m<sup>2</sup> |
| 3. Ligand-receptor affinity constant at zero load $K_a^0$ | 0.069 μm<sup>2</sup> |
| 4. Distance between particles and endothelial cell layer at equilibrium $\Delta$ | 45 nm |
| 5. Ligand-receptor bond length $\lambda$ | 5 nm |
| 6. Drag coefficient for spherical particles $F^s$ | 1.668 |

**Figure 4:**
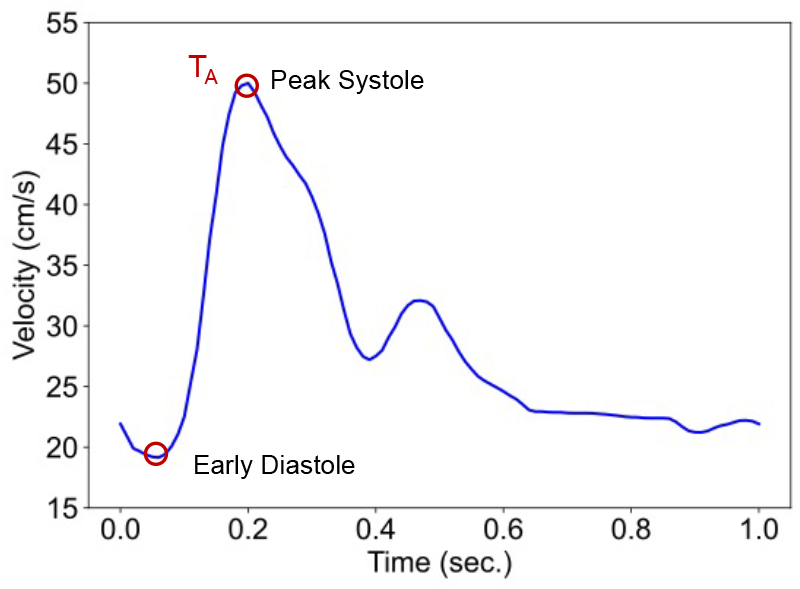
Velocity waveform in a healthy carotid artery. Time points *T*_*A*_ and *T*_*B*_ correspond to the maximum and minimum inflow velocities, respectively.

### 3.1 Experimental validation of the numerical method

Carefully designed microfluidic chamber experiments were carried out to validate the proposed method. A 3D microfluidic channel of 5*×* 5 *×*30 mm was employed in which endothelial cells that were activated with tumor necrosis factor alpha (TNF-*α*) to upregulate vascular adhesion molecule (VCAM) expression were attached to the bottom wall. Nanoparticles of poly(lactide-co-glycolide)/hyaluronic acid block copolymer (PLGA-*b*-HA) conjugated with VHSPNKK peptide, and having varying diameters, were injected into the chamber and particles that attached to the C166 mouse endothelium were imaged. The chamber was then exposed to wash flows with particle-free media at different flow rates that are observed in typical arteries, and the particles that retained on the endothelium were imaged again. This data was used in model validation. For numerical simulations, the base fluid was chosen as cell culture media, which is an incompressible Newtonian fluid and consistent with what was used in the experiments. A constant parabolic inflow velocity corresponding to a given, but otherwise arbitrary Reynolds number (Re) was prescribed at the inlet wall while stress free boundary conditions were applied at the outlet. No slip velocity boundary conditions were applied on all other walls. Nanoparticles with average diameter of 220 nm and 750 nm were introduced into the chamber in a time-lagged manner. The process consisted of two stages: a nanoparticle injection stage which was followed by a wash stage with particle-free media. During the nanoparticle injection stage, a 2 mL solution with a nanoparticle concentration of 0.8 mg/mL was injected into the chamber at a flow rate corresponding to the specified Reynolds number. For 750 nm particles, the initial concentration of nanoparticle solution during the injection stage was kept at 0.16 mg/mL. Attachment of nanoparticles to the inflamed endothelial cell layer was facilitated by particle-cell adhesion model via Robin boundary conditions during the injection stage. In the subsequent wash stage, a 6 mL particle-free fluid was passed through the chamber to study the detachment of the bound nanoparticles. In this stage, the proposed asperity and dispersion models were also active in the boundary layer. While particles were injected at Re = 1 for all the test cases, washing was done both at Re = 1 and Re = 5 to study the effect of flow conditions (found in arteries) on retention of the bound particles.

Simulation results matched closely with the *in vitro* experimental data as shown in Figure 5. For comparison of different sets of data, concentration values at the center of bottom channel wall were normalized w.r.t the concentration of the injected nanoparticles solution. These results validate the numerical model that can accurately predict nanoparticles’ adhesion and retention under varying flow conditions, particle sizes, and polymer types.

**Figure 5:**
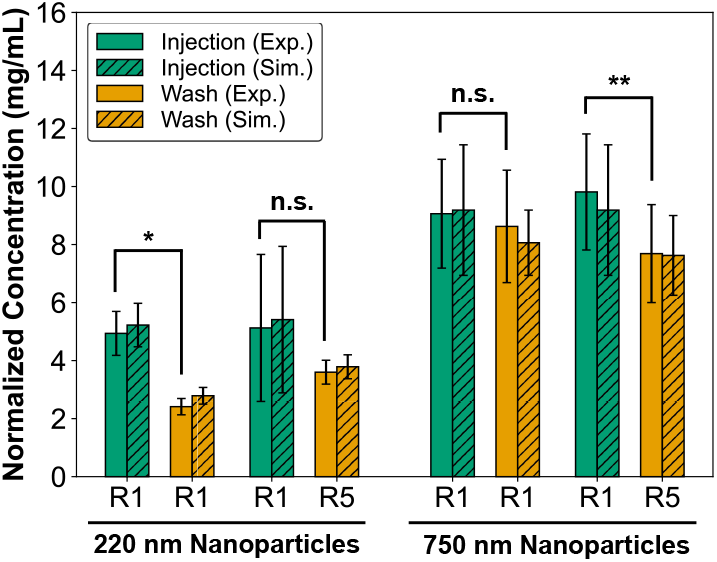
Comparative analysis of the adhesion and retention of nanoparticles between numerical simulations and *in vitro* experiments. The concentration of nanoparticles at the center of the blood vessel-simulating channel bottom was quantified at the end of nanoparticle injection (for adhesion) and after the wash flow with particle-free media (for retention). Particle concentration was normalized w.r.t the concentration of the injected nanoparticle solution during the injection stage. The proposed model was validated for PLGA-*b*-HA-VHSPNKK particles with an average diameters of 220 nm and 750 nm. R1 and R5 on the x-axis represent Reynolds numbers 1 and 5 of the media flow, respectively. The error bars represent a standard error in each dataset. n.s., *, and ** indicate not statistically significant with p *>* 0.05, p *<* 0.05, and p *<* 0.005, respectively.

**Figure 6:**
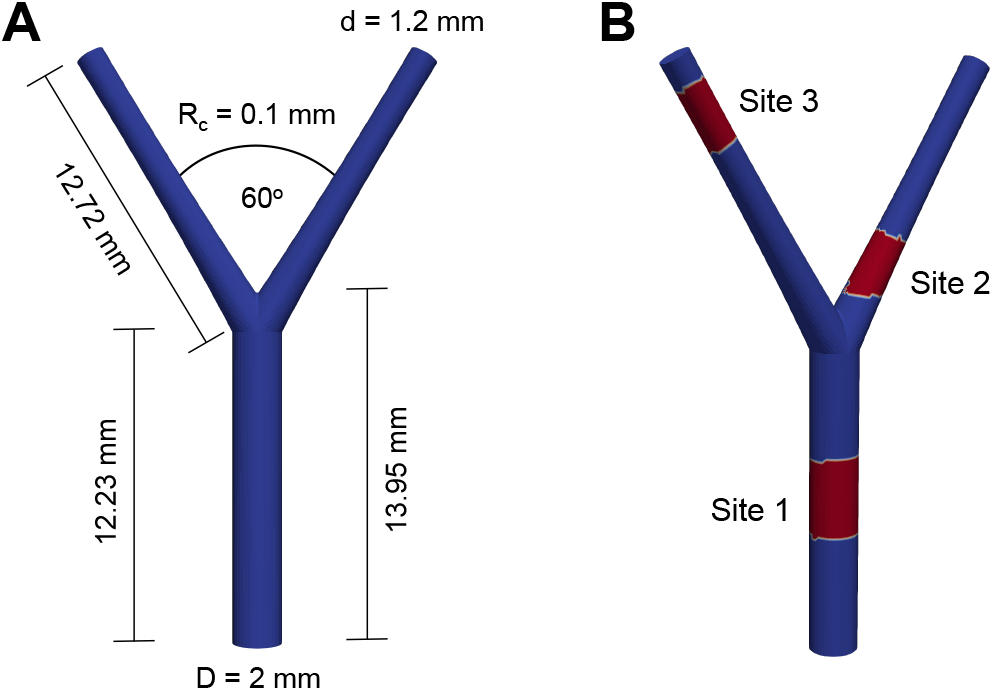
Three-dimensional idealized bifurcating artery. (A) Schematic diagram of the the idealized bifurcating artery. (B) Three targeted sites, highlighted in red, for drug delivery.

### 3.2 Idealized bifurcating artery

The first test case is a three-dimensional engineered bifurcating artery proposed in [52]. It consists of a parent artery where a pulsatile inflow (Figure 4) corresponding to the flow in a healthy carotid artery is prescribed. The velocity magnitudes are adjusted to match the average velocity that is found in typical smaller arteries or arterioles. The parent artery is 2 mm in diameter, 12.23 mm in length up to the bifurcation, and 13.95 mm in length to the bifurcation apex. The daughter arteries are both 1.2 mm in diameter and 12.72 mm in length from the bifurcation. The radius of curvature (*R*_*c*_) is 0.1 mm, and the angle between the daughter artery walls is 60 ^*°*^. No-slip boundary conditions are prescribed at the outer walls of the geometry. Stress-free outflow boundary conditions are applied at the end of the daughter arteries. The discretization consists of 33,883 nodes and 21,802 quadratic tetrahedral elements. The time step chosen is 0.01 s.

For targeted drug delivery, three random sites are specified on the geometry. Site 1 is in the parent artery, while site 2 and site 3 are located near the bifurcation and near the outlet on the two daughter arteries, respectively. The particle-cell adhesion model, asperity model, and dispersion model are active in the boundary layers of these targeted sites. Nanoparticles, characterized by an average diameter of 220 nm and a concentration of 0.1 mg/mL in blood, are introduced into the system at the inlet over a duration of 30 cardiac cycles. Subsequently a wash phase ensues during which blood flow that is devoid of nanoparticles is passed through the geometry for an additional 30 cycles. The wash cycle helps in assessing the extent of drug retention at the targeted sites against the dislodging hydrodynamic forces.

Figure 7 presents velocity, shear rate, viscosity, and pressure profiles in the bifurcating artery at two distinct time points: *T*_*A*_ representing peak systole and *T*_*B*_ corresponding to early diastole. At early diastole, the flow fields exhibit significantly reduced values in comparison to peak systole, primarily because this phase coincides with the minimum velocity magnitude on the velocity waveform (Figure 4). Additionally, higher flow velocities occur near the outlets of the two branching arteries due to their smaller diameters, resulting in maximum shear rates along the outer walls adjacent to these outlets. In contrast, the bulk flow at the artery’s center exhibits minimal shear rates due to lower velocity gradients, resulting in the highest viscosity in the bulk flow and a decrease in the boundary layer. This viscosity variation follows the shear-thinning blood flow model, with viscosities nearing *µ*_∞_ during peak systole and returning closer to the initial viscosity *µ*_0_ during early diastole. At the artery’s inlet, pressure is at its peak and it gradually decreases along the length of artery. The laminar flow ensures there is no significant pressure spike at the bifurcation apex.

**Figure 7:**
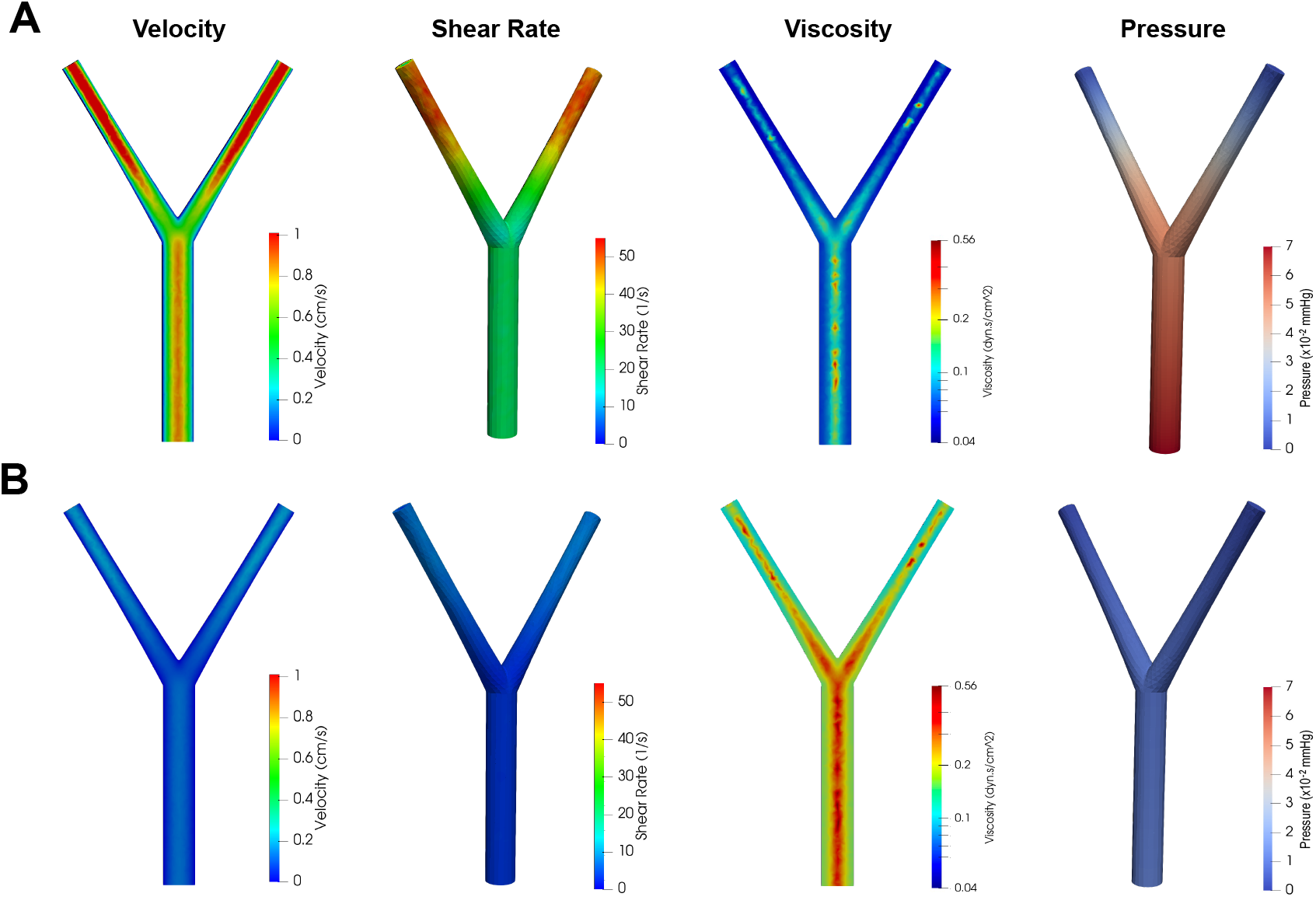
Various flow fields in idealized bifurcating artery at (A) peak systole (time point *T*_*A*_) and (B) early diastole (time point *T*_*B*_).

Effect of circadian rhythm on nanoparticle adhesion and retention is studied by simulating injection and wash stages during sleep and wake cycles (Figure 7). Circadian variation in physiological parameters such as inflow velocity, blood pressure, and heart rate are determined from the experimental data presented in Figure 2 where magenta and yellow lines represent sleep and wake cycles, respectively. Wake cycle has 25% higher inflow velocity, 20% greater pressure, and 27% faster heart rate as compared to the sleep cycle. Since each of the injection and wash stages spans 30 cardiac cycles, a greater heart rate in the wake cycle translates to lesser injection and wash duration. At the end of the injection stage, the concentration of attached particles at target site 1 does not differ significantly between sleep and wake cycles. However, at sites 2 and 3, particle adhesion is 11% and 25% higher, respectively, during the sleep cycle compared to the wake cycle. However, at the end of wash stage, all three target sites show significantly better drug retention in the sleep cycle as compared to the wake cycle. This is due to the higher pressure and flow rates during the wake cycle that induce stronger dislodging forces. Sites 1, 2 and 3 have 32%, 60%, and 59% higher average particle retention in the sleep cycle as compared to the wake cycle. Time evolution of retention of attached particles during the wash stage is shown in Figure 7D. It is evident that the concentration of attached particles during the wake cycles drop quicker, as greater number of particles get detached under stronger dislodging forces. After 25 seconds, concentration of particles that are retained at the target sites 1 and 2 during the sleep cycle is 23% and 18% higher than the wake cycle, respectively. The study provides evidence that the circadian variation in physiological parameters significantly affect particle retention at the target sites in the blood flow. This test case highlights that the circadian rhythm can be exploited in improving retention time of the nanoparticles and, in turn, improving the efficacy of drug delivery.

To better understand how the location of the target site within an arterial branch impacts particle adhesion and retention, the time evolution of the mean concentration of particles at sites 1 and 2 is plotted in Figure 8 for both sleep and wake cycles. Numerical simulations indicate that the concentration at site 1, located in the parent artery, plateaus at 0.34 mg/mL during the injection stage. Any further particle deposition is washed away as the adhesion capacity of the site is maximized. In contrast, the concentration at site 2, located near the bifurcation, continues to increase. At bifurcations, blood flow creates regions of disturbed flow that favor nanoparticle accumulation. The formation of low-velocity regions on the inner side of the branch in this case increases the retention of nanoparticles near the vessel walls, promoting their adherence to the target endothelium layer as evidenced in the case of target site 2. During the wash stage, the concentration of particles at the sites decreases as the external flow dislodges some of the attached particles. This behavior is consistent across both wake and sleep cycles. The simulation showed that variations in vessel geometry also affect the hydrodynamic forces experienced by nanoparticles. Optimized particle design can exploit these forces for enhanced targeting and retention in specific regions.

**Figure 8:**
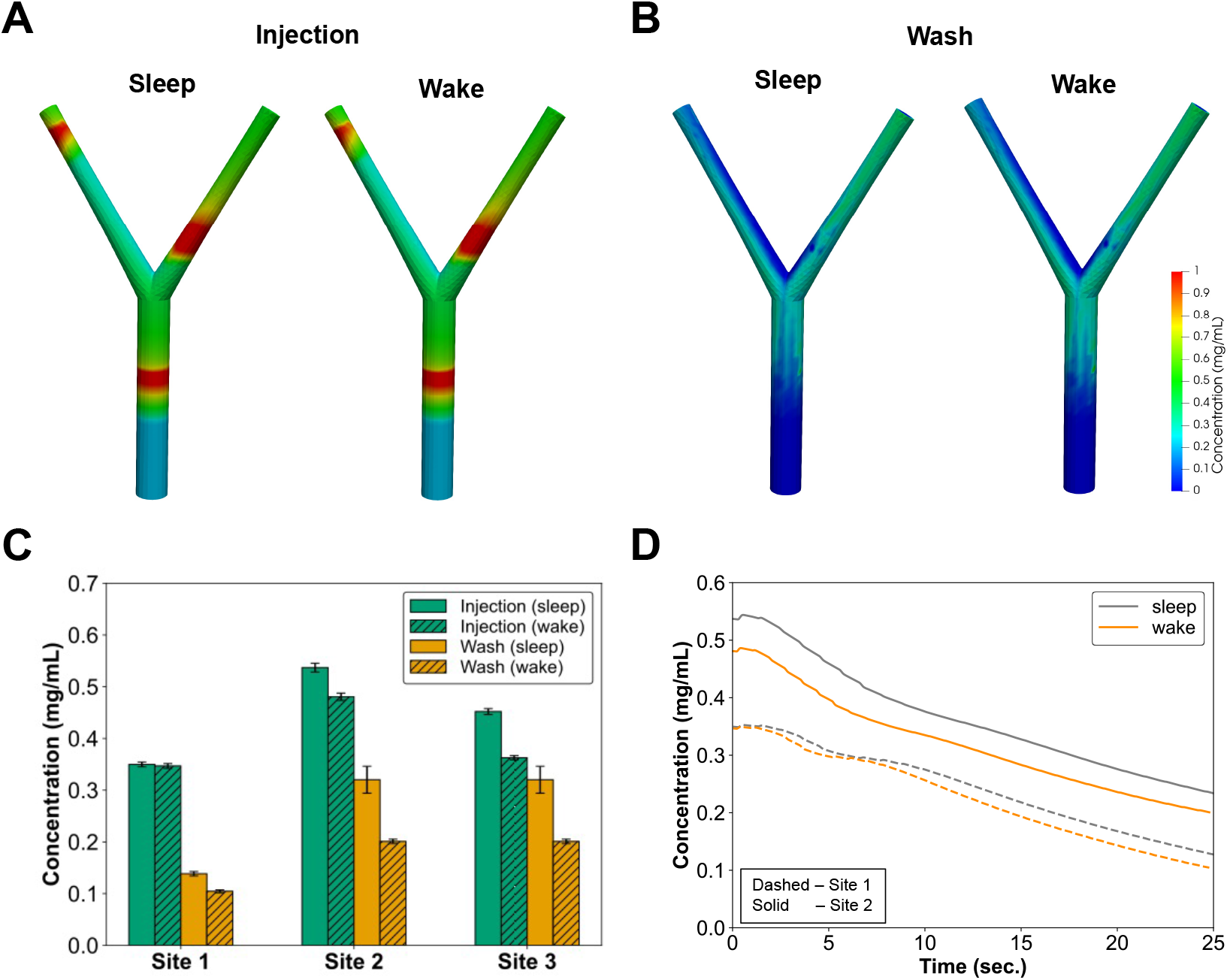
Effect of circadian rhythm on drug adhesion and retention in an idealized bifurcating artery. Particle adhesion at the three target sites at the end of (A) injection stage, and (B) wash stage is qualitatively shown. (C) Comparison of mean concentration across the three target sites at the end of injection and wash stages. Particle adhesion at sites 2 and 3 is 11% and 25% more in the sleep cycle as compared to the wake cycle, respectively. While negligible variation is observed between the two cycles at site 1, all three target sites show better drug retention in the sleep cycle as compared to the wake cycle. (D) Evolution of drug retention during wash stage for sleep and wake cycles. After 25 seconds of wash flow, the concentrations of drug particles at sites 1 and 2 are 23% and 18% higher, respectively, during the sleep cycle as compared to the wake cycle

**Figure 9:**
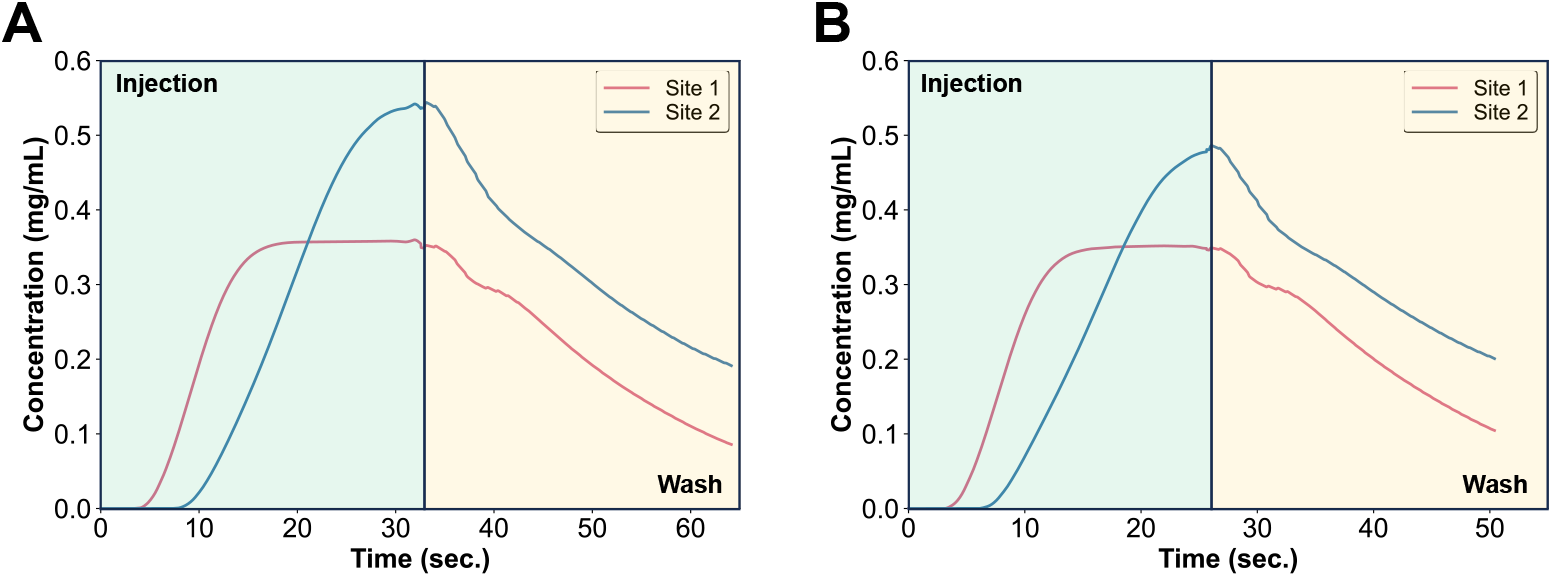
Time evolution of nanoparticle concentration at the target sites 1 and 2 during (A) sleep cycle and (B) wake cycle.

### 3.3 Idealized arterial tree

Carotid artery stenosis restricts oxygen flow to the brain, which requires a constant oxygen supply for proper functioning. Even a brief interruption in blood and oxygen supply can lead to brain cell damage, that can get triggered within minutes. Severe narrowing or blockage of carotid arteries can result in strokes, and if plaque fragments break off, they can also obstruct brain blood flow, leading to strokes. In this test case, an idealized arterial system with a stenosis branch is used to simulate drug transport at the targeted stenosis sites.

The geometry of the arterial system, as shown in Figure 10, is proposed in [53]. The diameter at the inlet is 1.26 mm and it symmetrically bifurcates into two daughter branches with diameter 0.98 mm each. The bifurcation angle is 60^*°*^. After the first bifurcation, the two daughter branches with a diameter *D*_*p*_ asymmetrically bifurcate into a large branch with diameter 0.911*D*_*p*_, and a small branch with diameter 0.584*D*_*p*_. From the second bifurcation onwards, the bifurcation angle is set equal to 60^*°*^. Each branch keeps the same diameter (i.e., *D*_*i*_) along a length, 20*D*_*i*_. The healthy branch (i.e., the left branch in Figure 10) is constructed by following the above geometric ratios. For the stenosed branch (i.e., the right branch in Figure 10), we assume that the smaller downstream arteries are blocked at the bifurcations. Consequently, only large branches are constructed, while maintaining the same change in the diameters at the corresponding bifurcation locations as in the healthy branch. The drug delivery targets are precisely these blocked smaller arteries at the bifurcations, as indicated in Figure 10. Table 3 presents the diameters as well as lengths of all the branches of the full geometry shown in Figure 10.

**Table 3:** Diameters and lengths of branches in idealized arterial system.

| Label | Diameter (mm) | Length (mm) |
| --- | --- | --- |
| a | 1.26 | 25.2 |
| b | 0.98 | 19.6 |
| c | 0.572 | 11.44 |
| d | 0.334 | 6.67 |
| e | 0.195 | 3.89 |
| f | 0.304 | 6.08 |
| g | 0.521 | 10.42 |
| h | 0.475 | 9.5 |
| i | 0.277 | 5.54 |
| j | 0.433 | 8.66 |
| k | 0.893 | 17.86 |
| l | 0.814 | 16.28 |
| m | 0.742 | 14.84 |
| n | 0.253 | 5.05 |
| o | 0.394 | 7.89 |
| p | 0.676 | 13.52 |
| q | 0.616 | 12.32 |

**Figure 10:**
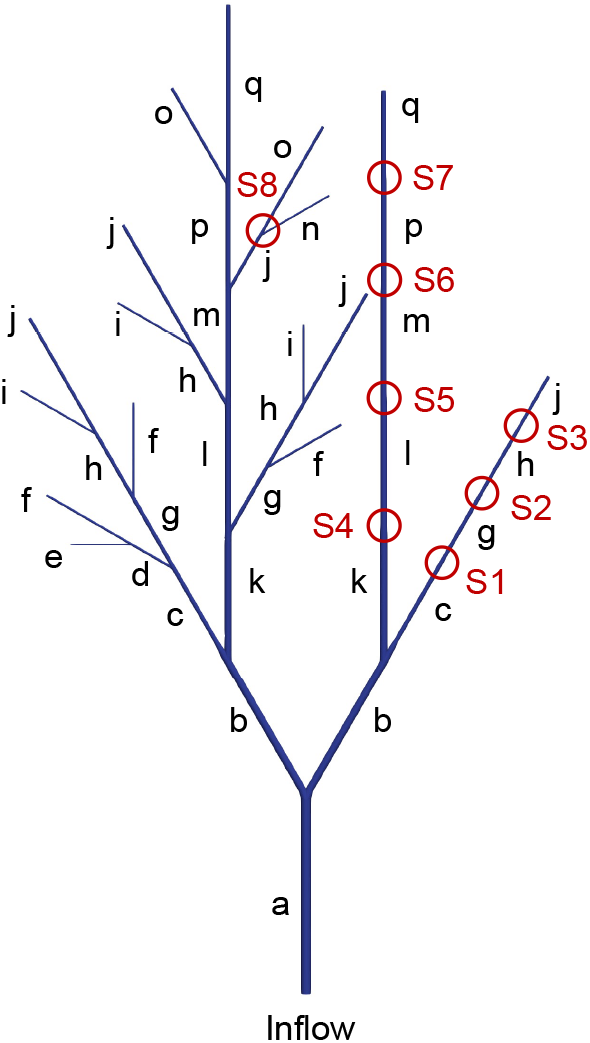
Idealized arterial system geometry with an inlet at the bottom and multiple outlets. Left side represents healthy branch with further bifurcations into smaller arteries while the right side represents a stenosed branch with blocked smaller arteries at the bifurcations. Thus, only large branches are constructed on the right side. The drug delivery targets (S1 to S7) are these blocked bifurcations. An additional site S8 in the left branch is activated halfway through the injection stage to simulate particle adhesion to target endothelial cells appearing at bifurcations. The geometric details of each branch is given in Table 3.

The geometry is discretized into 217,365 quadratic tetrahedral elements with 127,144 nodes. The problem is run with a time step of 0.01 s. A parabolic velocity profile is prescribed at the inflow with an average velocity of 3 mm/s following the typical waveform as given in Figure 4. Stress-free velocity boundary conditions are applied at the outlets while no-slip velocity boundary conditions are applied at all other boundary walls. For the pressure field, resistance boundary conditions at the two outlets are applied such that the pressure at the outlet is 120 mmHg during systolic and 80 mmHg during diastole.

Figure 11A shows the velocity magnitude contours in the arterial tree. Since the stenosed branch has larger downstream resistance as compared to the healthy branch due to its blocked smaller arteries, the healthy branch (i.e., left bottom portion in Figure 11A) shows larger velocity magnitude than in the stenosed branch (i.e., right bottom portion in Figure 11A). A similar pattern is observed for shear rates that are higher in the upstream healthy branch as compared to the stenosed branch. As we go downstream in the healthy branch, the flow velocity reduces in the smaller arteries. However, the opposite behavior is observed in the stenosed branch where smaller arteries have higher shear rates due to the blocked bifurcations.

**Figure 11:**
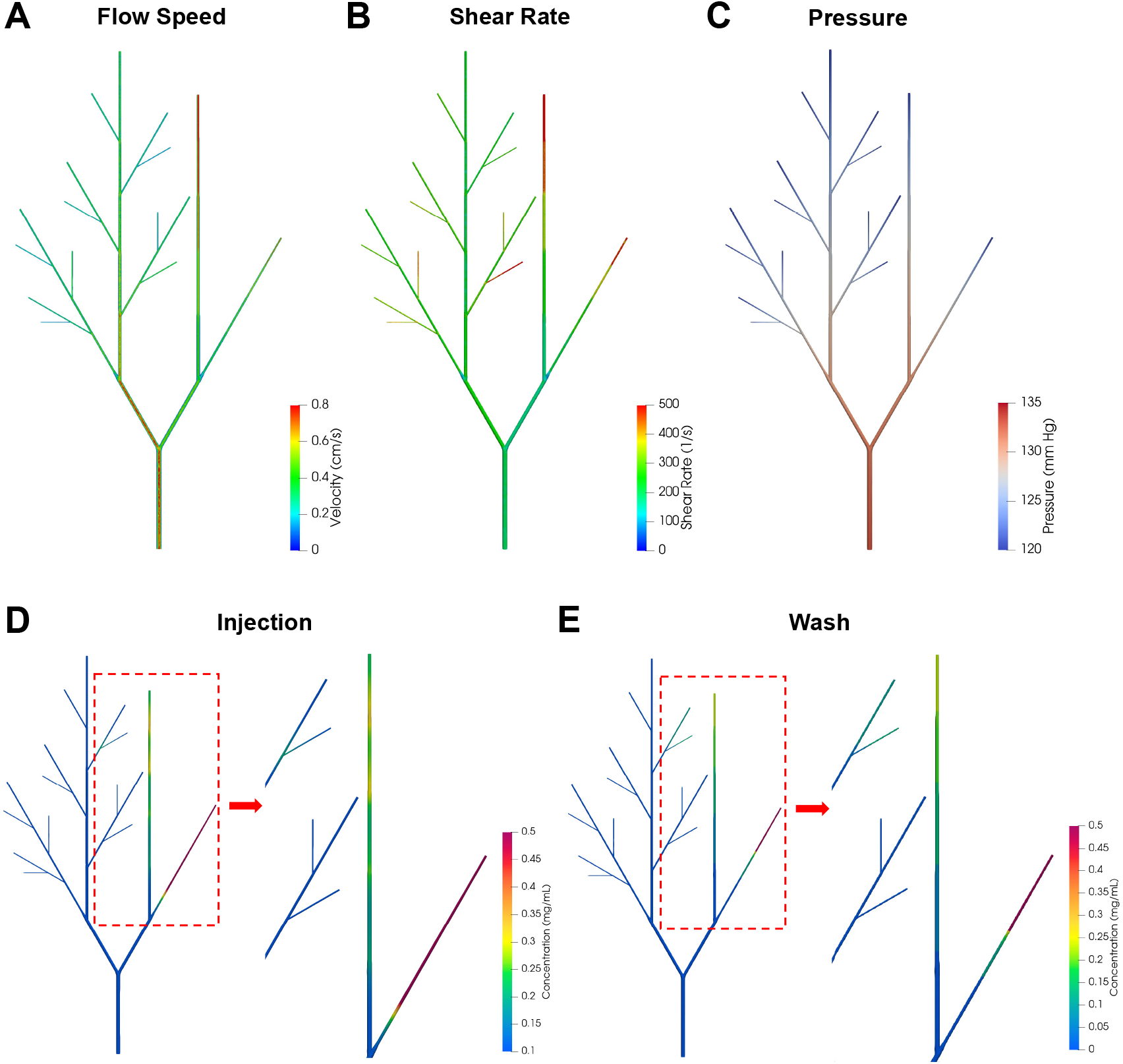
Spatiotemporal dynamics of particle transport, attachment, and retention in an idealized arterial tree with targeted stenosis sites. (A) Velocity magnitude contour at peak systole T_A_. (B) Shear rate at peak systole T_A_. (C) Pressure profile at early diastole T_B_. (D) Concentration profile at the end of injection stage, and (E) wash stage. Concentration build-up can be observed at the targeted sites (S1 to S7) in the stenosed branch as well at site S8 in the healthy left branch. The concentration of particles then drops down at the end of the wash stage.

In order to transport ligand-coated particles to the target stenosis sites, nanoparticles with an average diameter of 220 nm were injected into the bloodstream at the inlet using a solution with a concentration of 0.1 mg/mL. Attachment of nanoparticles to the stenosis sites was governed by particle-cell adhesion model via Robin boundary conditions. Figure 11D shows concentration profile of particles at the end of the injection stage. At arterial bifurcations, the blood flow exhibits local variations in flow velocity and shear stress, which are critical in determining endothelial cell behavior and function. These hemodynamic changes can lead to altered endothelial cell responses and are associated with regions that are prone to atherosclerosis due to low or oscillatory shear stresses [54]. To simulate this phenomenon, an additional target site S8 was introduced in the healthy left branch which gets activated halfway during the injection stage. Concentration buildup due to the attached particles is seen at all the target sites. In the stenosis branch, a bulk of drug goes into the sub-branch with target sites S1 to S3 resulting in the maximum concentration of attached particles at those sites (Figure 12). In the subsequent wash stage, a particle-free blood flow was maintained for another 30 cardiac cycles. Almost all of the particles were washed away at sites S1 and S4, while 12%, 30%, 26%, 48%, 62%, and 17% of the particles were retained at sites S2, S3, S5, S6, S7, and S8, respectively at the end of wash stage. Site 3 showed maximum adhesion and retention of nanoparticles.

**Figure 12:**
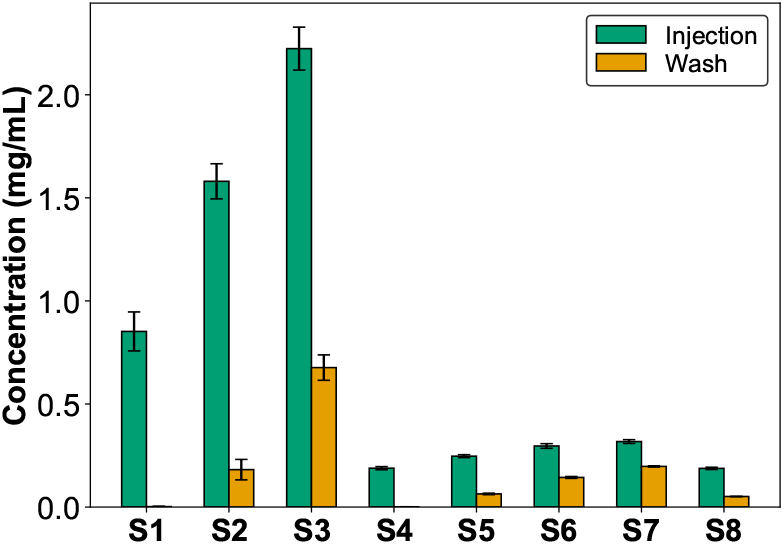
Comparison of mean concentration across all the target sites in the idealized arterial tree at the end of the injection and wash stages.

### 3.4 *In vivo* carotid artery system

In this test case, we apply the drug delivery computational framework with non-Newtonian blood flow model to *in vivo* geometric model obtained from MRI scans as shown in Figure 13. A computational grid is superimposed on a patient-specific 3D model of the carotid arterial system, consisting of 388,508 nodes and 243,839 quadratic hexahedral elements. The computational geometry includes the common carotid artery, which ascends within the neck and bifurcates into the internal and external carotid arteries. The two target sites, S1 and S2, are located in the ophthalmic artery and anterior cerebral artery, respectively, which branch off from the internal carotid artery. The anterior cerebral artery supplies blood to the medial portions of the primary motor and sensory cortices, and any blockage can lead to a stroke, affecting leg motor and sensory functions. Target site S3 is situated in the frontal branch of the superficial temporal artery, which supplies blood to the scalp and forehead tissue. The anatomy of the computational geometry along with the three target sites is illustrated in Figure 13.

**Figure 13:**
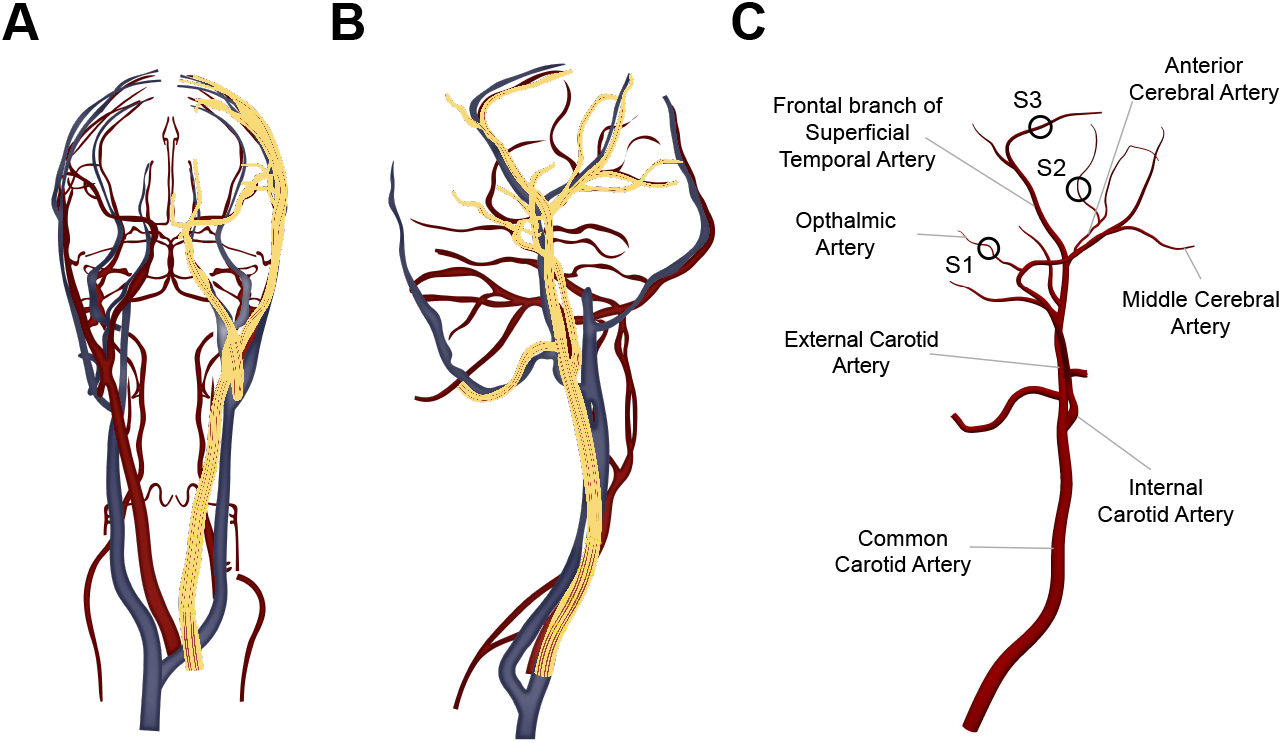
*In vivo* geometric model of arterial system in the brain along with the superposed computational grid (highlighted in yellow). (A) Frontal view (B) Side view (C) Anatomy of modeled geometry with target sites.

A physiologically relevant pulsatile inflow velocity of 35 cm/s is applied at the inlet located in the common carotid artery. No-slip velocity boundary conditions are applied at all the vessel walls. The resistance boundary condition in equation (15), which produces a physiological pressure waveform at the outlets, is imposed to incorporate the downstream resistive effects of the smaller branching arteries. The resistance parameter *R* for each branch is tuned based on the flow rate in the branches, calculated with traction-free boundary conditions, to obtain a pressure amplitude between 85–125 mmHg at the outlets. The simulation is run with a time step of 0.01 seconds. The coupled system of equations converges quadratically in the nonlinear iterative solution procedure at each time step. The pressure and velocity decrease sharply as blood flows into the smaller branches of the external and internal carotid arteries, as shown in Figure 14(A-C). The shear rates in the smaller branches where the target sites are also located range between 10-100 s^−1^, which is typical for microvasculature. Figure 14A presents a volume rendering of the magnitude of the velocity field. Figure 14(B-C) shows instantaneous snapshots of the shear rate and pressure field at peak systole, where these quantities reach their maximum value due to higher blood velocity. The anatomical features, including the curvature and geometry of blood vessels, influence various aspects of the blood flow, such as velocity profiles and shear stress distribution. Shear stress exhibits greater fluctuations at locations of curvature and bifurcations (Figure 14B), which in turn influences drug adhesion and retention.

**Figure 14:**
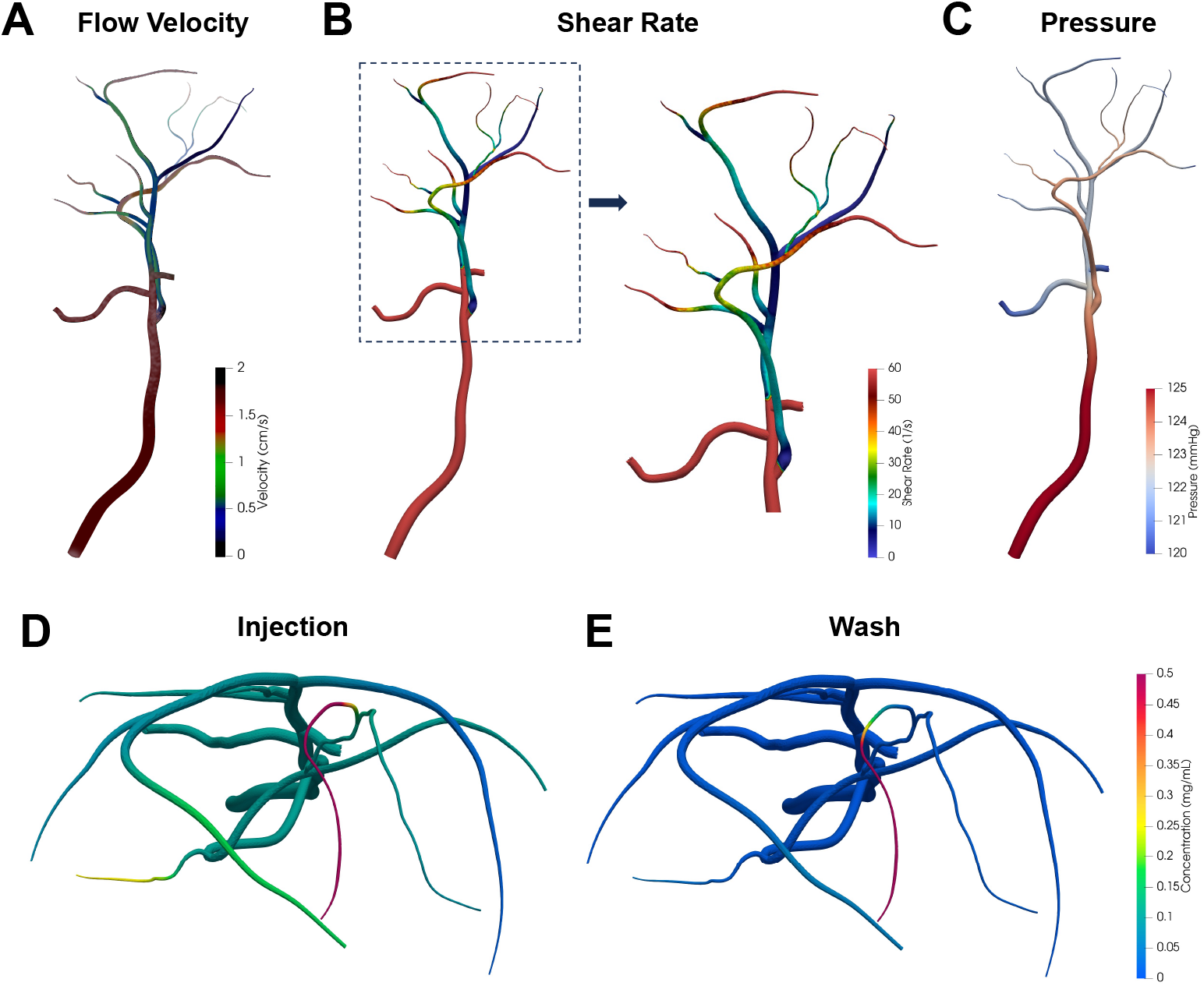
Flow physics in the carotid artery system at peak systole. (A) flow velocity, (B) shear rate, and (C) pressure. (D) Nanoparticles that are injected at the inflow get attached at all the target sites as shown by the buildup of nanoparticle concentration at the end of the injection stage. The maximum buildup is in the anterior cerebral artery. (E) A fraction of the attached particles gets dislodged under the effect of hydrodynamic forces during the wash stage.

For drug delivery, nanoparticles are injected at the inlet for 60 cardiac cycles. The particles are transported to the target sites by the flow of blood, where they attach under the influence of particle-cell adhesion forces. Figure 15(A-B) shows the time evolution of the mean concentration at the three target sites. Adhesion and retention of particles at the anterior cerebral artery (site 2) are an order of magnitude higher compared to the other two sites located in the ophthalmic artery and the frontal branch of the superficial temporal artery. This highlights the significant impact of the local geometric shape as well as the location of the target site on particle adhesion and retention. Site 2, located at a curvature of the blood vessels, experiences non-uniform shear stress distribution, with areas of lower shear stress typically found on the inner wall of the curve. These areas provide a favorable environment for nanoparticle accumulation due to reduced shear forces acting on the particles, allowing them to remain in contact with the vessel wall for longer periods [55]. Adhesion at site 1 saturates after 40 seconds, resulting in a plateau in the time evolution plot (Figure 15A). Site 2 shows an approximately linear increase in the concentration of attached particles, while site 3 shows an exponential increase in particle concentration. After the injection, wash stage ensues for another 60 cardiac cycles. During the wash stage, the concentration drop is most rapid at site 1. At the end of the wash stage, 36%, 43%, and 48% of particles are retained at target sites 1, 2, and 3, respectively (Figure 15C). The simulation tool developed in this paper help in understanding the impact of vessel curvature and geometry on patient-specific *in vivo* models with geometric uncertainties, allowing for the strategic design of nanoparticles that preferentially accumulate in target areas, ensuring localized and controlled drug release.

**Figure 15:**
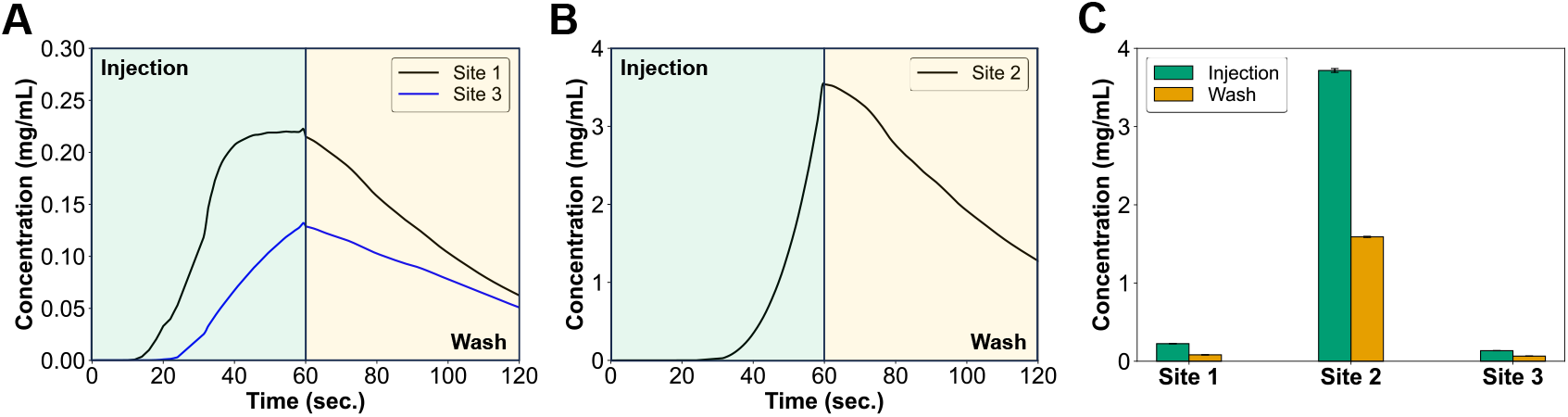
Time evolution of mean concentration at (A) target sites 1 and 3, as well as (B) target site 2, during the injection and wash stages. Adhesion and retention of particles at the anterior cerebral artery (site 2) is an order of magnitude higher as compared to the other two sites. (C) Comparison of mean concentration across all the target sites at the end of injection and wash stages. At the end of wash stage, 36%, 43%, and 48% particles are retained at target sites 1, 2, and 3, respectively.

## 4. Conclusions

We have presented a computational model to investigate nanoparticle-based targeted drug delivery under the influence of circadian rhythm-modulated blood flow in virtual *in vivo* arterial geometries. The primary objective was to understand how complex arterial geometries and physiological changes over the circadian cycle affect the transport, adhesion, and retention of nanoparticles within the vasculature. Our computational framework integrates a shear-rate-dependent non-Newtonian blood flow model with drug adhesion models, accounting for the complexities of nanoparticle transport in realistic arterial geometries that are derived from MRI scans. The simulations incorporated various factors influencing nanoparticle behavior, including vessel curvature, bifurcations, and downstream resistance, to provide a comprehensive understanding of the drug delivery dynamics. The results demonstrate that vessel geometry and location significantly impact nanoparticle adhesion and retention. Curved regions and bifurcations, characterized by non-uniform shear stress distributions, create favorable conditions for nanoparticle accumulation, highlighting the importance of the target site selection for optimized drug delivery. Circadian variations in physiological parameters, such as blood pressure and flow velocity, were shown to influence nanoparticle retention. During the sleep cycle, lower shear forces and flow rates resulted in higher nanoparticle retention as compared to that during the wake cycle. This finding underscores the potential of chronotherapy to enhance drug delivery efficacy by aligning treatment schedules with the body’s natural circadian rhythms. Our computational framework, validated through *in vitro* experimentation, serves as a virtual platform that can provide comprehensive insights into the variation of hemodynamic parameters across patient-specific *in vivo* models in response to physiological changes in blood flow and vessel geometries. These simulations also show that the local variations significantly affect particle adhesion and retention at the target sites. By leveraging these insights, we can not only enhance the design of nanoparticles that preferentially accumulate in target areas, but also optimize the timing of the therapeutic interventions. This computational modeling approach ultimately improves the effectiveness and safety of targeted drug delivery protocols.

## Acknowledgements

This research was supported by NIH-USA Grant No. R01GM135921. Computing resources for the simulations were provided by the TeraGrid/ACCESS Program under NSF Grant TG-DMS100004.

## Conflict of Interest Statement

The authors declare no potential conflict of interest.

## Notes

### Competing Interest Statement

The authors have declared no competing interest.

### Summary of Updates

Figure 2 has been updated and moved to Figure 1 in the new manuscript.

